# Separate neural dynamics underlying the acquisition of different auditory category structures

**DOI:** 10.1101/2021.01.25.428107

**Authors:** Gangyi Feng, Zhenzhong Gan, Han Gyol Yi, Shawn W. Ell, Casey L. Roark, Suiping Wang, Patrick C. M. Wong, Bharath Chandrasekaran

## Abstract

Current models of auditory category learning argue for a rigid specialization of hierarchically organized regions that are fine-tuned to extracting and mapping acoustic dimensions to categories. We test a competing hypothesis: the neural dynamics of emerging auditory representations are driven by category structures and learning strategies. We designed a category learning experiment where two groups of learners learned novel auditory categories with identical dimensions but differing category structures: rule-based (RB) and information-integration (II) based categories. Despite similar learning accuracies, strategies and cortico-striatal systems processing feedback differed across structures. Emergent neural representations of category information within an auditory frontotemporal pathway exclusively for the II learning task. In contrast, the RB task yielded neural representations within distributed regions involved in cognitive control that emerged at different time-points of learning. Our results demonstrate that learners’ neural systems are flexible and show distinct spatiotemporal patterns that are not dimension-specific but reflect underlying category structures.

**Significance:** Whether it is an alarm signifying danger or the characteristics of background noise, humans are capable of rapid auditory learning. Extant models posit that novel auditory representations emerge in the superior temporal gyrus, a region specialized for extracting behaviorally relevant auditory dimensions and transformed onto decisions via the dorsal auditory stream. Using a computational cognitive neuroscience approach, we offer an alternative viewpoint: emergent auditory representations are highly flexible, showing distinct spatial and temporal trajectories that reflect different category structures.

## Introduction

Learning to map continuous acoustic information to meaningful, behaviorally relevant auditory categories is critical to making sense of complex soundscapes. Behaviorally relevant sounds like speech and music contain multiple acoustic dimensions that are perceived differentially depending on listening experiences. Listeners need to extract relevant dimensions from continuously varying sound input and map the extracted structure to behaviorally relevant equivalence classes, or categories (Diehl et al., 2004). The successful acquisition of auditory category mapping requires our brain to efficiently reorganize to encode category-relevant acoustic signals for the formation of task-relevant neural representations and categorization decisions (Ashby and Maddox, 2005; Golestani and Zatorre, 2004; Ohl and Scheich, 2005; Schultz et al., 1998; Tricomi et al., 2006). Despite decades of behavioral, computational, and neuroimaging work examining auditory categorization, the neural dynamics underlying auditory category learning are poorly understood. Our objective here is to examine the neural dynamics in representations of two types of auditory category structures with identical dimensions: a rule-based (RB) category structure that is hypothesized to involve effortful, hypothesis-driven sound-to-rule mapping (Ashby et al., 1998), and an information-integration (II) category structure, that is hypothesized to involve a pre-decisional procedural-based integration of dimensions (Ashby et al., 1998; Ashby and Gott, 1988). These category structures are ubiquitous in the auditory world: speech sounds categories, for example, require pre-decisional integration of multiple dimensions that is difficult to verbalize and can be learned without explicit attention (Lim et al., 2019; Yi et al., 2016). In contrast, listeners also attribute complex rules to disambiguate sound categories (e.g., a physician listening to auscultation sounds).

One view of category learning posits a two-stage hierarchical model that does not differentiate between the learning of different category structures (Freedman et al., 2003; Jiang et al., 2007; Jiang et al., 2018). Specifically, the current two-stage models involve a perceptual learning phase in the sensory cortex (e.g., auditory cortex), followed by a higher-level learning phase in the prefrontal cortex, which can learn to identify, discriminate, or categorize stimuli. Per the two-stage model, auditory category learning involves increased selectivity in the representation of relevant auditory features within the auditory cortex (e.g., sharpening neuronal populations in auditory areas to form a task-independent reduction of the sensory input), which in turn serves as the input to prefrontal regions that implement categorical decisions (Jiang et al., 2018; Myers et al., 2009). The prefrontal cortex is not considered as the site of category sound representation *per se* but play important role in facilitating the formation of representations in the auditory cortex during learning (Myers, 2014). The frontal-temporal regions are structurally and functionally connected via dorsal and ventral auditory streams (Sheppard et al., 2012; Wong et al., 2011), providing the substrate for the sound-to-category mapping.

In contrast to the two-stage hierarchical model, the Dual Learning System (DLS) model (Chandrasekaran et al., 2014a; Chandrasekaran et al., 2014b) posits that distinct cortico-striatal learning systems drive different forms of auditory category learning. The neurobiology mechanisms proposed in the DLS model are derived from theoretical frameworks in vision (Ashby et al., 1998; Ashby and Maddox, 2005; Helie et al., 2015; Seger, 2008; Seger and Miller, 2010). Emerging literature examine multiple or dual learning systems across various domains, including automaticity (Ashby and Crossley, 2012; Ashby and Maddox, 2005, 2011), visual category learning (Carpenter et al., 2016; Nomura et al., 2007; Nomura and Reber, 2012; Seger, 2008; Seger and Miller, 2010) and speech category learning (Feng et al., 2019; Myers, 2014; Myers and Swan, 2012; Perrachione et al., 2011; Yi et al., 2016). In auditory domains, the DLS model proposes two distinct and parallel cortico-striatal circuits that underlie category learning: an explicit sound-to-rule stream that involves a cortico-thalamic-basal ganglia network composed of the prefrontal cortex (PFC), hippocampus, thalamus, head of the caudate nucleus, and anterior cingulate cortex (ACC), and a procedural-based sound-to-reward stream that involves the auditory-motor network in coordination with the reward-based striatal circuitry. Per the DLS model, during sound-to-rule mapping, a rule is generated within the auditory network and maintained in the prefrontal cortex (PFC) and hippocampus, and PFC and thalamus. The thalamocortical loop receives inhibitory projections from the globus pallidus (GP), which in turn receives inhibitory projections from the caudate. Excitation of the caudate by the PFC results in excitation of the PFC by the thalamus, providing a neural mechanism through which external corrective feedback on decisions can further maintain the pre-existing rule. In contrast, the sound-to-reward system implicitly maps sound to behavioral responses via the reward-sensitive striatal circuitry. This is achieved via many-to-one corticostriatal projections from the superior temporal cortex to the putamen. A closed-loop projection back to the STG allows the basal ganglia to fine-tune STG function. The sound-to-reward system is not consciously penetrable, and associates perception with motor actions that lead to reinforcements via feedback. Mechanistically, during sound-to-reward mapping, a single medium-spiny neuron in the putamen implicitly associates an abstract motor response with a cluster of sensory cells. Cortical-striatal synaptic learning is facilitated by a reinforcement signal (to correct feedback) from the ventral striatum (nucleus accumbens, NAC). The DLS model predicts that when the auditory category structure is discernable by rules, the *sound-to-rule* system dominates learning; however, when the category structure requires implicit integration across multiple dimensions, the *sound-to-reward* system is more optimal and may take over learning to ensure high categorization accuracies.

Here we generated novel non-speech auditory category structures with the same underlying dimensions, spectral and temporal modulation. These dimensions are considered to be the building blocks of complex signals like speech and music, where they are robustly represented in the auditory cortex (Schonwiesner and Zatorre, 2009). We used the same dimensions to create two category structures: a rule-based (RB) category structure allows categorization using an explicit decisional criterion on the spectral modulation dimension (high vs. low) and another on the temporal modulation dimension (fast vs. slow), and an information-integration (II) category structure that was created by rotating the separable structure by 45° clockwise. Optimal learning of the II category structure requires pre-decisional integration of spectral and temporal modulation dimensions to determine categories. Using non-speech auditory categories allows us to control for the learners’ prior auditory experiences with speech and music. Critically, it provides a flexible way to manipulate the category structures using the same dimensions while the behavioral learning performance can be comparable across the two category structures, overcoming the limitation in unbalanced performance or stimulus design in previous studies. We recruited 60 participants who were randomly assigned to either the II (*n* = 30) or RB (*n* = 30) groups in a between-subjects design. All participants in a group were actively trained on the same structure using a feedback-based sound-to-category training paradigm during fMRI scanning. We focused on examining group differences in sound-induced multivoxel pattern representations of category-relevant acoustic dimensions across training blocks (e.g., group-by-block interaction effect). Per the two-stage perceptual hierarchy model, irrespective of category structure (RB vs. II), we expect neural representations to emerge within superior temporal gyrus (STG) and inferior frontal gyrus (IFG) ensembles at different time-points in learning. In contrast, per the DLS model, we predicted that representations are task and category-structure contingent such that representations emerge within the auditory-motor circuitry for II category structures and within the distributed rule-based circuitry for the RB category structure.

## Materials and Methods

### Participants

Young adult native Mandarin speakers (*n* = 60; mean age: 21.2 years; SD = 2.1; age range: 18-28 years; all right-handed) with no hearing problems (based on a self-report questionnaire) were recruited from the South China Normal University community and received monetary compensation for their participation. Participants were excluded if they self-reported any major psychiatric conditions, neurological disorders, hearing disorders, head trauma, or use of psychoactive drugs or psychotropic medication. All participants had a normal or corrected-to-normal vision and minimal formal music training experience (< 1 year). Participants were randomly assigned to either the RB or II group (*n* = 30 each). The two groups of participants were matched in age (*t*_(58)_ = 0.249, *P* = 0.805), gender (II: 22 females; RB: 19 females), and years in musical training (*t*_(58)_ = 1.644, *P* = 0.106). The recruitment, consent, testing, and compensation procedures were approved by the South China Normal University Institutional Review Board and The Joint Chinese University of Hong Kong – New Territories East Cluster Clinical Research Ethics Committee (The Joint CUHK-NTEC CREC).

### Experimental design and Stimuli

Consistent with (Yi and Chandrasekaran, 2016), 40 sounds per category structure were generated by modulating a white noise stimulus (duration = 0.5 sec; digital sampling rate = 44.1 kHz; low-pass filtered at 4.8 kHz). Spectral modulation frequency ranged from 0.25 to 1.33 cyc/oct. Temporal modulation frequency ranged from 4 to 10 Hz (Fig. 1A). These modulation frequencies were selected because they are strongly represented in the human auditory cortex (Schonwiesner and Zatorre, 2009). The amplitude of modulation depth was 30 dB. All sounds were RMS amplitude normalized to 80 dB. To create the RB category structure, 40 coordinates were first generated in an abstract normalized two-dimensional space, with a minimum value of 0 and the maximum value of 1. Four bivariate normal distributions of 40 coordinates were centered on (0.33, 0.33), (0.33, 0.68), (0.68, 0.33), and (0.68, 0.68), with a standard deviation of 0.1 for both dimensions. Values along each dimension were logarithmically mapped onto spectral and temporal modulation frequencies. Thereby, for the RB category structure, the optimal decision boundaries reflect the placement of a decision criterion along the spectral modulation dimension at 0.58 cyc/oct and along the temporal modulation dimension at 6.32 Hz (Fig. 1A right panel, dashed lines). The II category structure was created by rotating the RB category structure by 45^°^ counterclockwise. The optimal decision boundaries for the II category structure are not easy to describe verbally and rely on both spectral and temporal modulation dimensions (Fig. 1A left panel, dashed lines).

**Fig. 1.**
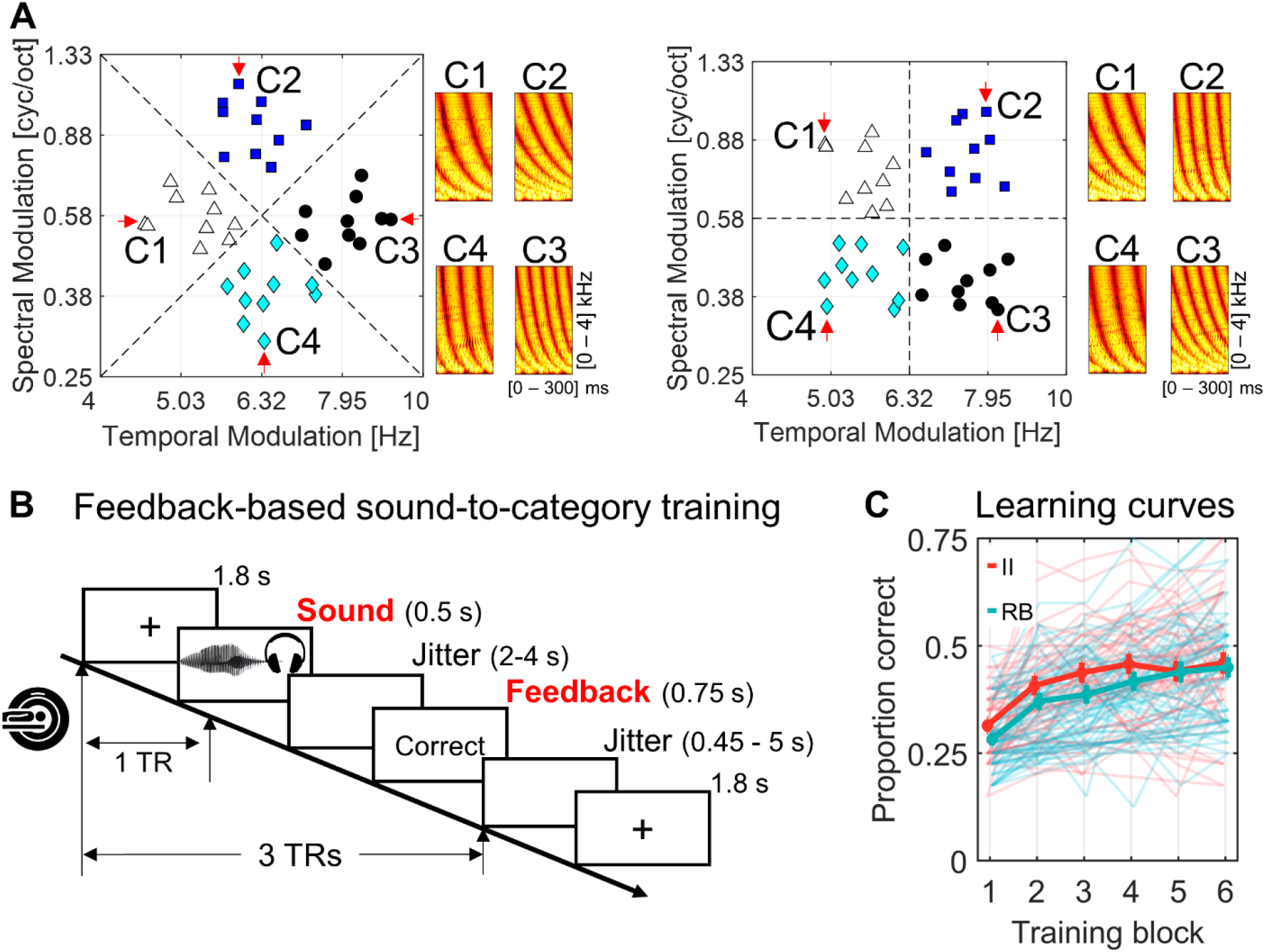
Auditory dimensions, category structures, training paradigm, and behavioral learning performance. **A**, two category structures and spectrograms of sample stimuli. Left panel: forty sounds from the II structure. Four categories (C1 - 4) of sounds from the II structure were plotted in different shapes/colors in a two-dimensional (i.e., spectral and temporal modulation dimensions) perceptual space. Right panel: forty sounds from the RB structure were plotted in the same perceptual space. Dashed lines reflect optimal decision boundaries between the categories. **B**, the feedback-based sound-to-category training procedure used in the fMRI experiment. TR = scanning repetition. Each training trial consisted of three TRs. Each sound was presented in a silence gap (0.8 sec; sounds duration = 0.5 sec) of a TR. **C**, categorization accuracies increased over training blocks comparably for both groups. No significant difference was found between groups across blocks.

### Sound-to-Category Training Procedures

Sounds were presented via MRI-compatible circumaural headphones in an MR scanner (Siemens 3T Tim Trio MRI system). Visual stimuli (e.g., feedback) were presented using an in-scanner projector visible using a mirror attached to the head coil. Participants were equipped with a 2-button response box in each hand. The experiment consisted of 6 contiguous scan runs or “training blocks.” Before each block, participants were instructed to attend to the fixation cross on the screen (10 sec). On each trial, a sound category was presented during the 800-ms silence gap following the 1700-ms image acquisition (Fig. 1B; also see Imaging acquisition section for the customized sparse-sampling imaging parameters). Participants were instructed to categorize the sounds into one of four categories. Informational corrective feedback (“正确, 这是类别 4。” [“RIGHT, that was a 4.”] or “错误, 这是类别 4。” [“WRONG, that was a 4.”]) was displayed for 750 ms after each response. If the participant failed to respond within a two-sec window, cautionary feedback was presented (“ 没反应” [“NO RESPONSE”]). After feedback presentation, a fixation cross was displayed until the onset of the next imaging acquisition. Each trial lasted three TRs (7.5 sec in total). Single null trials (i.e., silence trial; *n* = 10 per block) with a fixation cross were randomly inserted between sound trials to jitter the inter-trial intervals for better estimation of single-trial activations. To separate stimulus-response and feedback-evoked neural responses, the stimulus-to-feedback interval was pooled from random samples from a uniform distribution of 2 to 4 sec. Each stimulus was presented once within a block. The presentation order of the stimuli was randomized for all participants and blocks.

### Imaging acquisition

MRI data were acquired using a Siemens 3T Tim Trio MRI system with a 12-channel head coil in the Brain Imaging Center at South China Normal University. Functional images were acquired using a sparse-sampling T2^*^-weighted gradient echo-planar imaging (EPI) pulse sequence [repetition time (TR) = 2,500 ms with 800-ms silence gap, TE = 30 ms, flip angle = 90^°^, 31 slices, field of view = 224 × 224 mm^2^, in-plane resolution = 3.5 × 3.5 mm^2^, slice thickness = 3.5 mm with 1.1 mm gap]. T1-weighted high-resolution structural images were acquired using a magnetization prepared rapid acquisition gradient echo sequence (176 slices, TR = 1,900 ms, TE = 2.53 ms, flip angle = 9^°^, voxel size = 1 × 1 × 1 mm^3^).

## Statistical analyses

### Behavioral data modeling

Participants’ trial-by-trial accuracy and reaction times were analyzed using linear mixed-effects (LME) regression analysis to assess the main effect of group and training block. The fixed effects of interest were group (II = 0 and RB = 1) and the number of the blocks (1-6) mean-centered to 0. Analysis of variance for the LME models was conducted to reveal the statistical significance for the fixed factors and the interaction effect.

Besides, we used a behavioral representational similarity analysis (bRSA) to evaluate the emerging representation of category-relevant perceptual information by modeling learners’ block-by-block categorization responses. We first converted participants’ categorization responses into a response confusion matrix (a 40 × 40 pair-wise matrix; same response = 1 and different responses = 0) for each block. We then correlated a predefined representational dissimilarity matrix (RDM) with the response confusion matrices using Spearman’s rank correlation. The predefined RDM was constructed by calculating the standardized Euclidean distance between each pair of sounds in the two-dimension (spectral and temporal modulation) perceptual space (see the II and RB RDMs in Fig. 2A). Because the sounds from both structures are drawn from the same distribution in a standardized perceptual space, the correlation between the II and RB RDMs is almost at the ceiling (*r* = 0.97, *P* < 0.001), as expected.

**Fig. 2.**
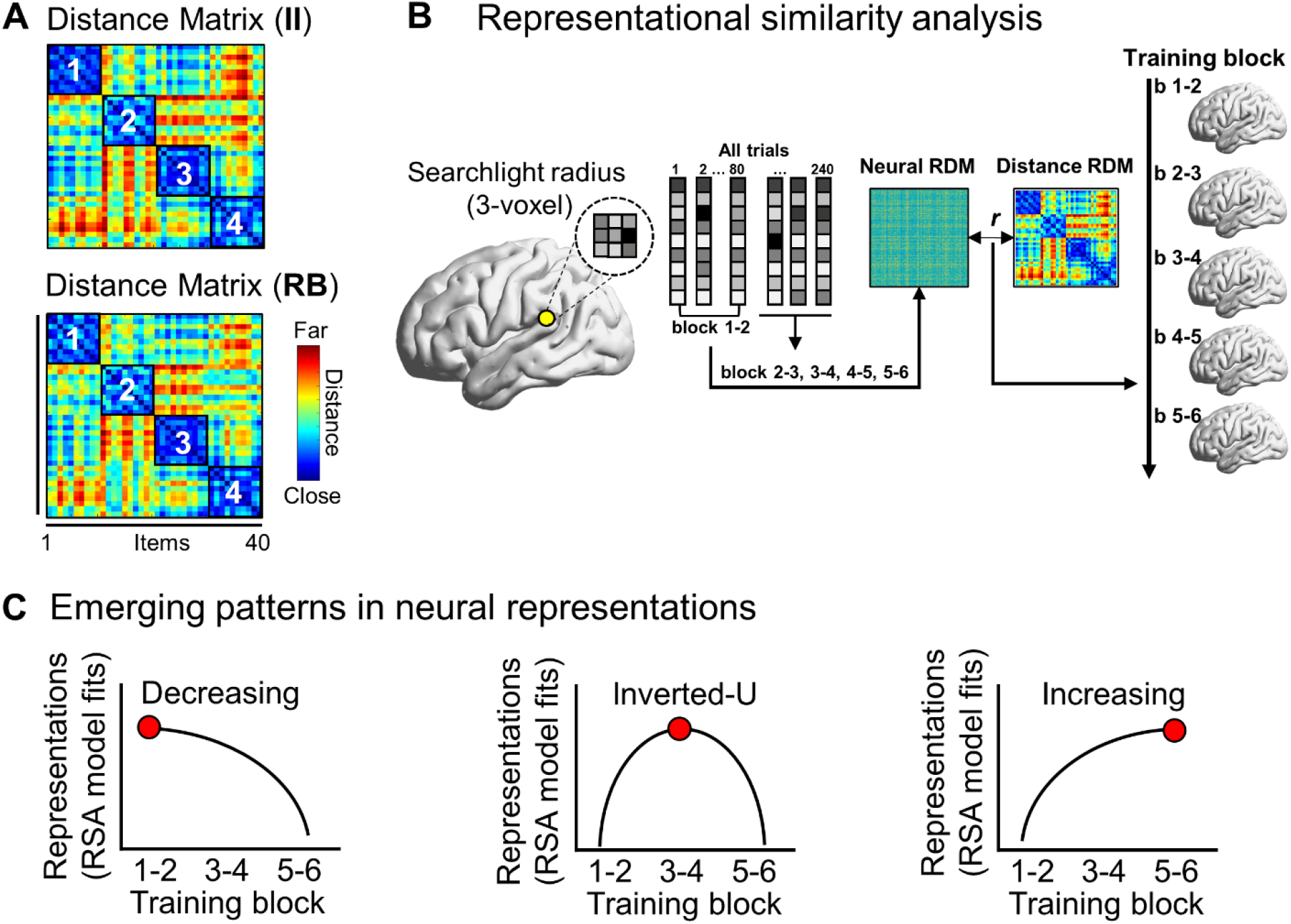
Multivariate representational similarity analysis (RSA) procedure and putative emerging patterns in neural representations. **A**, perceptual distance matrices (i.e., RDM) for II and RB stimulus structures. The RDMs were created based on pair-wise standardized Euclidean distance in a two-dimensional perceptual space. Within-category item pairs were labeled and highlighted. **B**, the searchlight-based RSA procedure. RSA was conducted across the whole brain by correlating the local neural RDMs with perceptual category distance RDMs for each group. Data from two consecutive blocks were combined (sliding window = 2 blocks). **C**, conceptual diagrams of the three hypothesized training-induced emerging patterns in neural representations of category information. The neural representations are assumed to be formed and updated following training. Three emerging patterns (decreasing, inverted-U, and increasing) are illustrated with line graphs. Red circles represent the most robust representations in a specific training phase.

### Model-Based Analyses

To get a more detailed description of how participants categorized the stimuli, a number of different decision-bound models (Ashby, 1992a; Maddox and Ashby, 1993) were fit separately to the data for each block and each learner. Decision bound models are derived from general recognition theory (Ashby and Townsend, 1986), a multivariate generalization of signal detection theory (Green and Swets, 1966). It is assumed that, on each trial, the percept can be represented as a point in a multidimensional psychological space and that each participant constructs a decision bound to partition the perceptual space into response regions. The participant determines which region the percept is in, and then makes the corresponding response. While this decision strategy is deterministic, decision bound models predict probabilistic responding because of trial-by-trial perceptual and criterial noise (Ashby and Lee, 1993). Below we briefly describe the decision bound models. For more details, see Ashby (1992a) or Maddox and Ashby (1993). The classification of these models as either *rule-based* or *information-integration* models is designed to reflect current theories of how these categories are learned (e.g., Ashby et al., 1998) and has received considerable empirical support (see Ashby and Valentin, 2017 for a review).

#### Rule-Based Models

##### Unidimensional Classifier (UC)

This model assumes that the stimulus space is partitioned into four regions by setting three criteria on one of the stimulus dimensions. Two versions of the UC were fit to these data. One version assumes that participants attended selectively to spectral modulation and the other version assumes participants attended selectively to temporal modulation. The UC has four free parameters, three correspond to the decision criteria on the attended dimension and the other corresponds to the variance of internal (perceptual and criterial) noise.

##### Conjunctive Classifier (CC)

A more appropriate rule-based strategy given the current category structures (Fig. 1A) is a conjunction rule involving separate decisions about the stimulus values on the two dimensions, with the response assignment based on the outcome of these two decisions (Ashby and Gott, 1988). All versions of the CC assume that the participant partitions the stimulus space into four regions (i.e., if low on temporal modulation and high on spectral modulation, respond 1; if high on temporal modulation and high on spectral modulation, respond 2; if low on temporal modulation and low on spectral modulation, respond 3; if high on temporal modulation and low on spectral modulation, respond 4). The CC has three free parameters: the decision criteria on the two dimensions and a common value of internal noise for the two dimensions. A special case of the CC, the *optimal rule-based classifier*, assumes that participants use the CC that maximizes accuracy (i.e., the dashed boundary plotted with the RB structure in Fig. 1A). This special case has one free parameter (internal noise).

##### Conjunctive* Classifier (CC*)

This class of models is similar to the CC with the exception that they assume two criteria on either temporal modulation or spectral modulation. The first CC* model assumes that the temporal modulation dimension is partitioned into three regions and that a criterion on spectral modulation is used for stimuli intermediate in temporal modulation, resulting in the following rule: respond 1 if temporal modulation is low; respond 4 if temporal modulation is high; respond 2 if temporal modulation is intermediate and spectral modulation is high; respond 3 if temporal modulation is intermediate and spectral modulation is low. The second CC* model assumes that the spectral modulation dimension is partitioned into three regions and that a criterion on temporal modulation is used for stimuli intermediate in spectral modulation, resulting in the following rule: respond 3 if spectral modulation is low; respond 2 if spectral modulation is high; respond 4 if spectral modulation is intermediate and temporal modulation is high; respond 1 if spectral modulation is intermediate and temporal modulation is low. The CC* model has four free parameters (two criteria on temporal/spectral modulation, one criterion on spectral/temporal modulation, and one for internal noise).

#### Information-Integration Models

##### The Linear Classifier (LC)

This model assumes that two linear decision boundaries partition the stimulus space into four regions (see the dashed boundary plotted with the information-integration structure in Fig. 1A for an example). The LC differs from the CC in that the LC does not assume decisional selective-attention (Ashby and Townsend, 1986). This produces an information-integration decision strategy because it requires linear integration of the perceived values on the stimulus dimensions prior to invoking any decision processes. The LC assumes two linear decision bounds of opposite slope (five parameters, slope, and intercept of both linear bounds and a common value of internal noise). A special case of the LC, the *optimal information-integration classifier*, assumes that participants use the LC that maximizes accuracy (i.e., the dashed boundary plotted with the II structure in Fig. 1A). This special case has one free parameter (perceptual and criterial noise).

##### The Minimum Distance Classifier (MDC)

This model assumes that there are four units (one associated with each category) representing a low-resolution map of the stimulus space (Ashby and Waldron, 1999; Ashby et al., 2001; Maddox et al., 2004). On each trial, the participant determines which unit is closest to the perceived stimulus and produces the associated response. Because the location of one of the units can be fixed, and because a uniform expansion or contraction of the space will not affect the location of the minimum-distance decision bounds, the MDC has six free parameters (five determining the location of the units and one for perceptual and criterial noise).

#### Random Responder Models

##### Equal Response Frequency (ERF)

This model assumes that participants randomly assign stimuli to the four categories in a manner that preserves the category base rates (i.e., 25% of the stimuli in each category). This model has no free parameters.

##### Biased Response Frequency (BRF)

This model assumes that participants randomly assign stimuli to the four categories in a manner that matches the participant’s categorization response frequencies. This model has three free parameters, the proportion of responses in categories 1, 2, and 3. Although the ERF and BRF are assumed to be consistent with guessing, these models would also likely provide the best account of participants that frequently shift to very different strategies.

#### Model Fitting

The model parameters were estimated using maximum likelihood (Ashby, 1992b; Wickens, 1982) and the goodness-of-fit statistic was

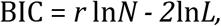

where *N* is the sample size, *r* is the number of free parameters, and *L* is the likelihood of the model given the data (Schwarz, 1978). The BIC statistic penalizes a model for poor fit and extra free parameters. To find the best model among a set of competitors, one simply computes a BIC value for each model, and then chooses the model with the smallest BIC.

## Neuroimaging data analyses

### Univariate activation analysis

All functional imaging data were preprocessed using SPM12 (Wellcome Department of Imaging Neuroscience, London, UK; www.fil.ion.ucl.ac.uk/spm/) closely following the pipeline described in previous studies (Feng et al., 2021; Feng et al., 2018; Feng et al., 2019). Here, we provide brief descriptions of these procedures. The T2*-weighted functional images were head-movement corrected. The high-resolution T1-weighted image was registered to the mean image of the functional images and further normalized to the Montreal Neurological Institute (MNI) space using a segmentation-normalization procedure for the estimation of normalization parameters. The realigned functional images were spatially smoothed using a Gaussian kernel (FWHM = 6 mm). Voxel-wise univariate activation analysis with the general linear model (GLM) was employed to examine the brain activity independently induced by sound categorization and feedback processing. For the subject-level analysis, a GLM with a design matrix including four regressors of interest (i.e., correct and incorrect sound categorization as well as correct and incorrect feedback presentations) was constructed for each participant. The regressors corresponding to the onset of trials were convolved with a canonical hemodynamic response function (without modeling temporal derivatives). Low-frequency drifts were removed using a temporal high-pass filter (cutoff at 128 sec). The AR(1) approach was used for autocorrelation correction. We identified outlier time points where the volumetric image was outside of three standard deviations (*SD*) of the global mean intensity or with the composite head movement that was larger than 1 mm (overall mean percentage of the outlier = 1.75 ± 1.05%; II learners = 1.93 ± 1.06%, RB learner = 1.57 ± 1.06%, two-sample *t*-test *t*_(58)_ = 1.32, *P* = 0.192). The outlier time points (1’s for outlier time points and 0’s for other time points) together with the six head-movement parameters and the session mean were added into the GLM models as nuisance regressors. The gray-matter image generated from the segmentation step was converted to a binary inclusive mask for each participant to define voxels of interest. For the group-level analysis, each statistical brain map was initially thresholded at voxel-wise *P* = 0.001. All reported brain regions were corrected at the cluster-level *P* = 0.05 using the family-wise error rate (FWER) approach as implemented in the SPM package.

### Representational similarity analysis (RSA)

For RSA, the realigned functional images in each participant’s native space (without spatial normalization) were analyzed with the subject-level GLMs to estimate single-trial brain responses using the least-squares single (LSS) approach (Mumford et al., 2014; Mumford et al., 2012). Specifically, a GLM with a design matrix for each trial was constructed and estimated separately. A design matrix consisted of a stimulus regressor of interest for that trial during the sound presentation; a regressor of non-interest consisted of all other events (i.e., feedback presentation for that trial, and sound and feedback presentation for the rest of the trials in the same block), outlier regressors, six head movement regressors, and a session mean regressor for each block individually. Therefore, 240 subject-level GLMs were constructed and estimated for each participant. The *t*-statistic brain maps were calculated for each trial and used for RSA (Misaki et al., 2010).

The RSA approach (Kriegeskorte and Kievit, 2013; Kriegeskorte et al., 2008) was used to examine the emerging neural representations of auditory category information as a function of sound-to-category training. The category information model as defined by the inter-sound perceptual distance (i.e., RDM; see Fig. 2A and the construction procedure in the *Behavioral data modeling* section) was used in the RSA. The model RDM was used to correlate the neural RDM (nRDM) derived from a spherical brain area (for the searchlight-based RSA) or a region of interest (for the ROI-based RSA) using Spearman’s rank correlation (see Fig. 2B for the graphical analysis procedure). Whole-brain RSA maps were calculated using the searchlight algorithm (Kriegeskorte et al., 2006) to identify brain regions that represent the category-relevant perceptual distance information during training. To increase the stability of the RSA estimation, single-trial data from two consecutive blocks (e.g., blocks 1 and 2, 2 and 3, etc.; see Fig. 2B, right panel) were combined to increase the signal-to-noise ratio (SNR) of sound item estimates by averaging the same items across two blocks. Therefore, five pairs of blocks were constructed. The multi-voxel activation patterns based on *t*-statistic values derived from each searchlight sphere (the average number of voxels in each sphere = 90) were used to calculate dissimilarities between each pair of sounds for the nRDMs (i.e, 1-Pearson’s correlation matrix). The nRDMs were then correlated with the predefined perceptual-distance RDM by using Spearman’s rank correlation. The correlation value of each sphere was standardized using Fisher’s *r*-to-*z* transformation and projected back to the center voxel to generate RSA maps. For group-level analysis, the searchlight RSA maps were first normalized to the MNI space and then fed to a flexible factorial-design analysis of variance (ANOVA) model as dependent variables to assess the main effects of group and training block as well as the interaction effect in multivariate representations. We also used a model-based approach to identify regions that fit different changing patterns of learning-related representational dynamics (see below for the description of three types of dynamic patterns). All group statistical maps from the multivariate analyses were thresholded at voxel-wise *P* = 0.005 with cluster-level FWER of 0.05.

We hypothesized that the neural representations of category information change as a function of the training block, with three types of dynamic changes in representations: increasing, decreasing, and inverted-U (see Fig. 2C for the diagram). These patterns assume that the neural representations of category-relevant information emerge at different time points of the training for different regions. This assumption has been supported by the observation of various learning studies that have demonstrated brain representations (mostly in univariate activations) of the learning stimuli change after training as compared with that of pre-training or early-training sessions, especially for the decreasing and increasing pattern (Karuza et al., 2014; Ley et al., 2012; Myers, 2014; Ohl and Scheich, 2005; Pasupathy and Miller, 2005; Tagarelli et al., 2019; Wang et al., 2003; Wong et al., 2007). The inverted-U pattern is motivated by learning studies interested in the function of the hippocampus memory system that show that the hippocampus temporally engages during learning to rapidly extract learning-relevant environmental information, mediate memory consolidation, and facilitate the transfer of new-acquired knowledge into the cortex (Baldassano et al., 2017; Davis and Gaskell, 2009; Schapiro et al., 2017; Takashima et al., 2014). Based on these observations, we hypothesized that the hippocampus memory system could temporally encode category-relevant information during learning. Therefore, an inverted-U pattern could reflect a pattern where the most robust representations emerge soon after training and a decrease in the following sessions. We are aware that the inverted-U pattern could range from several minutes to days of training depending on the nature of the learning task. Therefore, examining the inverted-U pattern of the representations in the current experiment is exploratory.

Each of these three patterns was proposed and modeled at the group-level analyses. The **increasing pattern** was defined as the neural representation increasing across training blocks (Fig. 2C, right panel). We predicted that this increasing pattern of the representation could closely follow the behavioral learning performance since the neural representations are hypothesized to subserve behavioral categorization. Therefore, an increasing curve function was created based on the group-level mean-centered behavioral learning accuracies (i.e., II = [−0.07, 0.00, 0.02, 0.02, 0.03]; RB = [−0.07, −0.02, 0.01, 0.03, 0.05]; each number denotes the weight of each pair of blocks [i.e., blocks 1-2, 2-3, 3-4, 4-5, and 5-6]). The **decreasing pattern** was defined as the neural representation decreasing across training blocks (Fig. 2C, left panel). The decreasing curve function was created by inverting the increasing function. The **inverted-U pattern** was defined as the neural representation emerging temporarily in the middle of training and diminishing in the “late” training phase (Fig. 2C, middle panel). The inverted-U function was created by assigning the highest weight for the “middle” blocks (e.g., block 3-4) while decreasing weights for the early and late blocks (i.e., [−0.8 0.2 1.2 0.2 −0.8]). The “middle” blocks could be arbitrarily defined. We also generated other variants of the inverted-U function to model the data, including assigning the highest weight to blocks 2-3 (i.e., [−0.8 1.2 0.4 0 −0.8]) and blocks 4-5 (i.e., [−0.8 0 0.4 1.2 −0.8]). These curve functions were used to weight and fit each pair of blocks’ RSA model correlations for each voxel across the whole brain, which can assess the extent to which a given region’s neural representation changes as a specific function.

## Results

### Behavioral response patterns and learning strategy modeling

Categorization accuracy significantly increased over training blocks for both groups (II learners: mean accuracy of the first block = 0.31 and last block = 0.46 [paired *t*-test: *t*_(29)_ = 5.38, *P* < 0.001]; RB learners: mean accuracy of the first block = 0.28 and last block = 0.45 [paired *t*-test: *t*_(29)_ = 6.60, *P* < 0.001]; Fig. 1C). The linear mixed-effects (LME) regression analysis (training block and group were two fixed factors and subject was a random factor) confirmed that the main effect of block in accuracy was significant (*F*_(5,348)_ = 26.64, *P* < 0.001) but the main effect of group was not significant (*F*_(1,348)_ = 1.43, *P* = 0.23). The behavioral representational similarity analysis (bRSA) further showed that the perceptual category distance RDMs were increasingly correlated with learners’ response confusion patterns for both groups (LME, main effect of block: *F*_(5,348)_ = 24.212, *P* < 0.001), but group differences were not significant (the main effect of group: *F*_(1,348)_ = 0.070, *P*_(1,348)_ = 0.792). No significant group-by-block interaction effect was found (*F*_(5,348)_ = 0.947, *P*_(1,348)_ = 0.451). These bRSA results demonstrate that learners’ response patterns were increasingly similar with the perceptual category distances between sounds for both groups.

We used computational modeling to assess each learner’s best-fitting strategy for each block. We coded the strategies into three categories, II, RB, and random responder (Fig. 3A). Participants who performed the II task (i.e., II learners) were more likely to use II strategies compared to other strategies (*χ*^2^ = 25.733, *df* = 2, *P* < 0.001). Similarly, participants who performed the RB task (i.e., RB learners) were more likely to use RB strategies (including unidimensional and conjunctive models) relative to other strategies (*χ*^2^ = 70.533, *df* = 2, *p* < 0.001). By using multinomial logistic regression analysis, we further confirmed that there were significant group differences (II vs. RB learners) in the proportions of the strategies used (II vs. RB strategy: *b* = 2.718, *SE* = 0.337, *t* = 8.067, *P* < 0.001; II vs. random strategy: *b* = 2.239, *SE* = 0.391, *t* = 6.391, *p* < 0.001; RB vs. random strategy: *b* = −0.478, *SE* = 0.268, *t* = −1.786, *P* = 0.074).

**Fig. 3.**
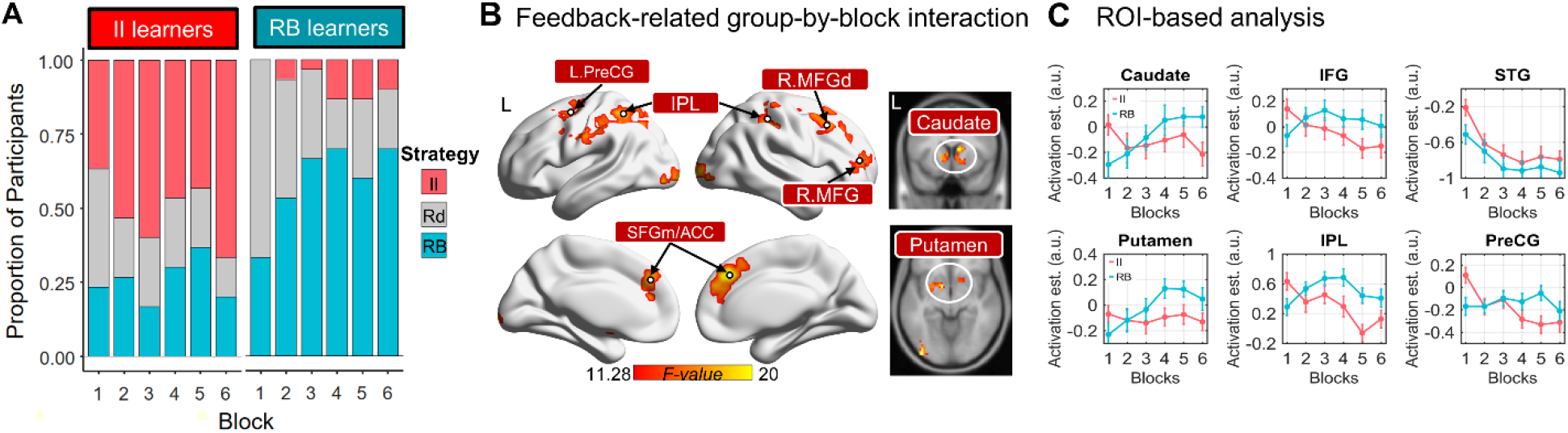
Learning strategy modeling results and group-by-block interaction effects in feedback-processing-related brain activations. **A**, the proportion of each learning strategy used in each block and learner group as revealed by neurocomputational decision-bound modeling. Strategy: II = optimal II or MDC responders; Rd = Random responders; RB = rule-based UC or CC responders. **B**, the group-by-block interaction effects were found in a distributed corticostriatal network. The brain map was thresholded at the voxel-level *P* = 0.001 and cluster-level FWER = 0.05. The frontoparietal and striatal regions were labeled. **C**, ROI-based analysis shows that the feedback-processing-related activations (feedback presentation vs. baseline) changed differently following training between the two groups in six anatomical-defined ROIs. Brain region abbreviation: PreCG, precentral gyrus; IPL, inferior parietal lobule; MFG, middle frontal gyrus; MFGd, dorsal middle frontal gyrus; SFGm, medial superior frontal gyrus; ACC, anterior cingulate cortex; IFG, inferior frontal gyrus; STG, superior temporal gyrus. L, left hemisphere; R, right hemisphere.

### Feedback-related corticostriatal activations changed differently between II and RB learners

We focus on examining the neural dynamics in feedback activations and sound representations elicited by training for each learning group and revealing group differences in the neural dynamics. Both univariate and multivariate neural measures (see Fig. 2A for the RSA procedure) were calculated and examined. For univariate activations, the feedback processing (i.e., feedback vs. baseline) yielded brain activations in the bilateral frontoparietal areas (bilateral inferior frontal gyrus [IFG]/middle frontal gyrus [MFG] and inferior parietal lobule [IPL]), insula, the left middle temporal gyrus (MTG), and occipital cortices (Fig. S1A, *Supplementary Information [SI]*). Although a larger extent of the frontoparietal areas was engaged for the RB learners than for the II learners, no region was found showing significant group differences (cluster-level FWER = 0.05). We also identified feedback-valence (i.e., correct vs. incorrect) related regions in the bilateral putamen and the head of the caudate nucleus where they showed greater activations for positive than negative (i.e., correct > incorrect) feedback for both groups (Fig. S1B), which consistent with previous findings in the context of speech category learning (Feng et al., 2019; Yi et al., 2016). Direct comparisons between II and RB learners in feedback-valence related activations (i.e., II _[correct-incorrect]_ vs. RB _[correct-incorrect]_) did not reveal any significant regions.

To examine the group differences in feedback-related activation across training blocks, we constructed a second-level flexible factorial-design analysis of variance (ANOVA) with the voxel-wise feedback-related activations (i.e., feedback vs. baseline and correct-incorrect feedback) as the dependent variable, group as a between-subject factor, and training block (i.e., block 1 to 6) as a within-subject factor (i.e., group-by-block two-way ANOVA). The learner group differences in dynamic changing patterns are summarized by the voxel-wise group-by-block interaction effect (Fig. 3B). A corticostriatal network was identified showing significant interaction effects, including the left precentral gyrus (L.PreCG), right ventral and dorsal MFG, bilateral IPL, bilateral medial superior frontal gyrus (mSFG)/anterior cingulate cortex (ACC), bilateral occipital cortex, bilateral putamen and head of caudate nucleus.

To further break down the group-by-block interaction effect, we performed ROI-based univariate-activation analyses in six pre-defined anatomical ROIs, including attention and cognitive-control-related frontoparietal regions (the bilateral IFG and IPL), striatum areas (the bilateral putamen and caudate nucleus) (related to reward and feedback processing), and sensorimotor regions (the bilateral STG and PreCG). These regions were selected because they have been proposed to be involved in category learning and representation of newly-acquired categories (Ashby and Maddox, 2005, 2011; Ashby and Valentin, 2017; Feng et al., 2018; Feng et al., 2019; Seger and Miller, 2010; Seger and Peterson, 2013; Yi et al., 2016). These ROIs were derived from the automated anatomical labeling atlas (AAL) (Rolls et al., 2015). The feedback-related activations for each block and subject were extracted and averaged across voxels within each ROI. The group-by-block activation profiles were showed in Fig. 3C. We did not find any ROI showing a significant main effect of group (*P*s > 0.1). In contrast, significant main effects of block were found in the bilateral STG (*F*_(5,348)_ = 9.41, *P* < 0.001), IPL (*F*_(5,348)_ = 4.90, *P* < 0.001) and PreCG (*F*_(5,348)_ = 3.58, *P* = 0.0036). Significant group-by-block interaction effects were found in the bilateral caudate nucleus (*F*_(5,348)_ = 3.58, *P* = 0.0036), putamen (*F*_(5,348)_ = 2.51, *P* = 0.029), IFG (*F*_(5,348)_ = 2.94, *P* = 0.013), IPL (*F*_(5,348)_ = 5.15, *P* < 0.001), and PreCG (*F*_(5,348)_ = 4.76, *P* < 0.001). These ROI-based findings are consistent with the whole-brain voxel-wise results shown in Fig. 3B and further reveal the changing dynamics of the feedback-related activations. These results further confirmed that a corticostriatal network, including the frontoparietal and striatum regions, were increasingly involved in the acquisition of RB categories, whereas the feedback-related activations in these regions decreased or remain stable during the learning of II categories.

### Group differences in emerging dynamics of the neural representations

The multivariate RSA was used to examine the emerging neural representations of category information (i.e., perceptual distance) for each group. We focused on analyzing the sound-categorization-related multivoxel patterns to reveal the extent to which the local activation patterns reflect the emerging neural representations. The whole-brain group-by-block analysis of variance (ANOVA) in neural representations of category distance information revealed a significant main effect of training block in the bilateral frontoparietal regions (Fig. 4A, the top panel), including the left lateral IFG and MFG, right IFG, bilateral posterior PreCG, and IPL, the supplementary motor area (SMA) and posterior cingulate cortex (PCC). The main effects of group were found in the left dorsal PreCG and left anterior IPL (Fig. 4A, the middle panel). Significantly more robust representations emerged for the II learners than that of the RB leaners in the two regions across training blocks (Fig. 4B, lower panel). Critically, we found another two regions, the left triangular IFG (L.IFGtri) and left STG (L.STG), that showed significant group-by-block interaction effects (Fig. 4A, bottom panel). The detailed interaction effects for the two regions are further displayed in the line graphs (Fig. 4B, upper panel), which shows that the representations increased following training for the II learners whereas the representations *decreased* for the RB learners.

**Fig. 4.**
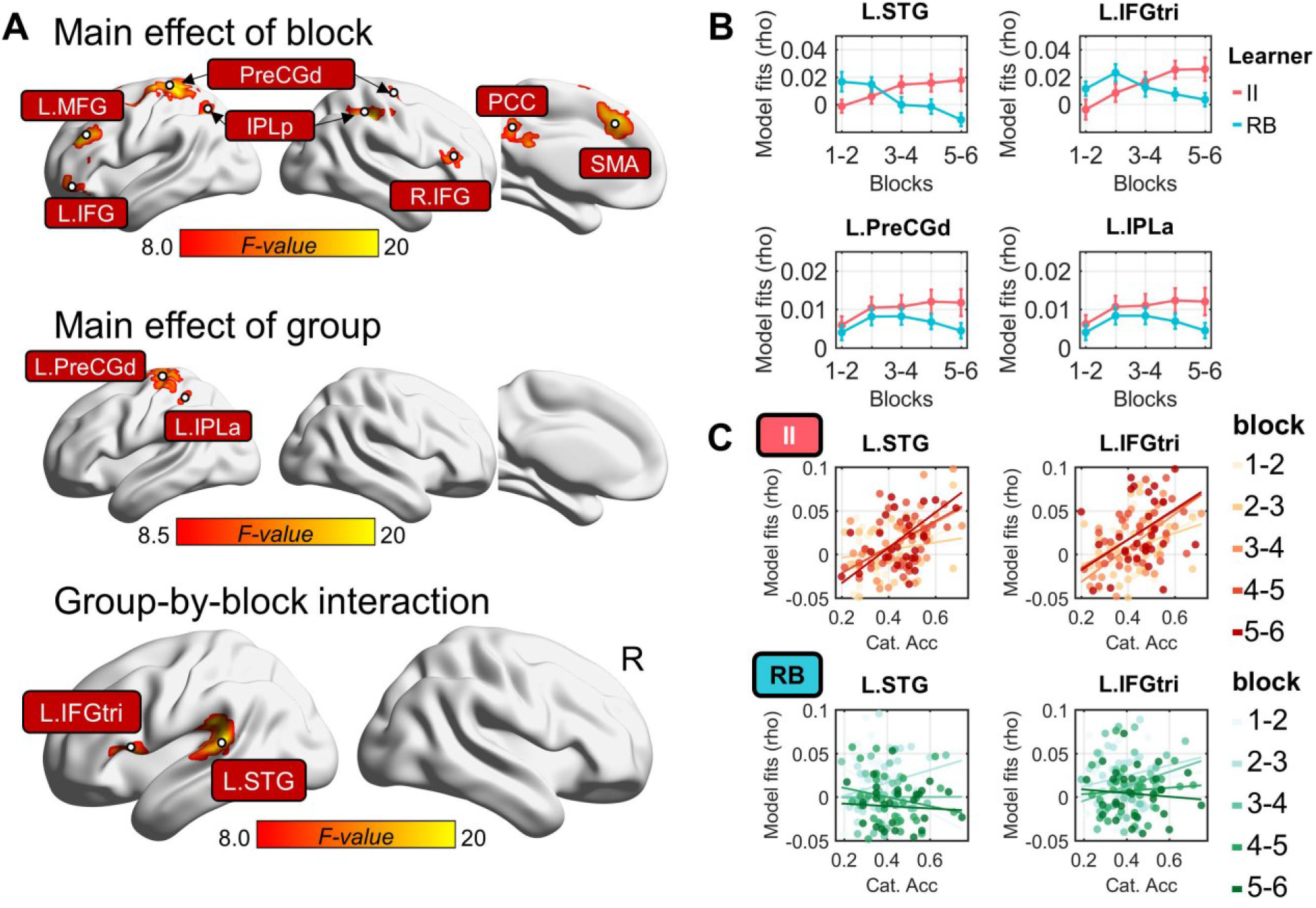
Whole-brain group-by-block ANOVA in the neural representations of category distance. **A**, the group-by-block ANOVA revealed the significant main effect of block (upper panel), the main effect of group (middle panel), and group-by-block interaction effects (bottom panel) in neural representations. Brain maps were thresholded at voxel-level *P* = 0.005 and cluster-level FWER = 0.05. **B**, changing profiles in neural representations for the two interaction-effect regions (L.STG and L.IFGtri) and the two main-effect-of-group regions (L.PreCGd and L.IPLa). These line graphs are for visualization purposes. **C**, individual differences in learning performance (i.e., categorization accuracy) were significantly correlated with the robustness of the emerging neural representations across blocks exclusively for the II learners. The lightness of dots and lines denotes training blocks.

To quantitatively compare the robustness of the emerging pattern between the two groups with a model-free approach that does not assume a priori the curve function across training blocks, we used a linear regression model to estimate the ‘emerging rate’ (i.e., slope) of the representations for each group separately. The independent variable is block pairs (i.e., block 1-2, 2-3, 3-4, etc.). The regression slope derived from the regression model is an indicator of the emerging rate where the higher slope indicates faster emergence in neural representations. We found that the representations in the L.IFG (peak coordinates: x = −42, y = 26, z = 1; cluster size = 738 mm^3^) and L.STG (extending to the supramarginal gyrus; peak coordinates: x = −63, y = 34, z = 10; cluster size = 1,359 mm^3^) increased significantly faster (i.e., higher slope) for II learners than that of RB learners. This result converged with the ANOVA findings that show significant group-by-block interaction effects in the two regions.

To examine the extent to which the emerging neural representations in the L.IFGtri and L.STG are behaviorally-relevant, we conducted correlation analyses between the robustness of the neural representations and behavioral categorization accuracies across training blocks. We performed an ROI-based inter-individual correlation analysis to reveal the behavioral-neural association for each group separately. Individual differences in the robustness of neural representations were significantly correlated with individual learning success (i.e., categorization accuracies) among II learners in the L.STG (block 1-2 [*r* = 0.31, *P* = 0.09], block 3-4 [*r* = 0.52, *P* = 0.003], block 5-6 [*r* = 0.54, *P* = 0.002]) as well as in the L.IFGtri (block 1-2 [*r* = 0.27, *P* = 0.14], block 3-4 [*r* = 0.62, *P* < 0.001], block 5-6 [*r* = 0.44, *P* = 0.015]) (Fig. 4C, upper panel), but not among RB learners (L.STG: block 1-2 [*r* = −0.26, *P* = 0.17], block 3-4 [*r* = 0.01, *P* = 0.99], block 5-6 [*r* = −0.06, *P* = 0.73]; L.IFGtri: block 1-2 [*r* = 0.11, *P* = 0.55], block 3-4 [*r* = 0.23, *P* = 0.21], block 5-6 [*r* = −0.09, *P* = 0.62]) (Fig. 4C, lower panel). The behavior-neural correlations for the II learners increase over training blocks (Fig. 4C, upper panel), which indicates that the learning performance is increasingly associated with the learners’ emerging neural representations. These results are consistent with the ANOVA results (see Fig. 4A&B).

### II learners: The emerging neural representations of category-perceptual distance

We hypothesized that the neural representations of category-related information change during training with different trajectories (Fig. 2B) for different learner groups. For the II leaners, univariate activation analyses revealed that sound categorization (compared with baseline) induced distributed fronto-temporoparietal activations, consistent with previous findings (Feng et al., 2019; Yi et al., 2016). This categorization network consists of the bilateral IFG, insula, PreCG and postcentral gyrus (PostCG), IPL, auditory cortices (e.g., bilateral Heschl’s gyrus [HG], and STG), and striatal areas (Fig. S2A, SI). Besides, we found a significant main effect of training block in a dorsal auditory pathway (Fig. S2B, SI), where sound categorization-related activations decreased following training, which may reflect stimulus-related repetition adaptation effects (Henson, 2003; Larsson and Smith, 2011; Summerfield et al., 2008). No region showed a significant main effect of group or group-by-block interaction effect.

The three hypothesized emerging patterns (i.e., increasing, decreasing, and inverted-U) were applied to reveal the neural representational changes across blocks. With the searchlight-based RSA and group-level weighted-contrast-based curve-fitting, we identified the neural representations of category-perceptual distance in a fronto-temporoparietal network showing a significantly increasing pattern mimicking the behavioral learning performance for the II learners (Fig. 5A). The most relevant regions involved were located in the bilateral IFG, left PreCG, left PostCG, left IPL, left supramarginal gyrus (L.SMG), left middle section of the superior temporal gyrus (L.STGm), right MFG, MTG, and PCC (see Table 1 for the region details). The increasing patterns were displayed in the line graphs for four representative regions (Fig. 5B). For visual comparison purposes, the dynamic changing patterns derived from the RB learners were also displayed for the same regions. No region showed a significant decreasing or inverted-U pattern.

**Fig. 5.**
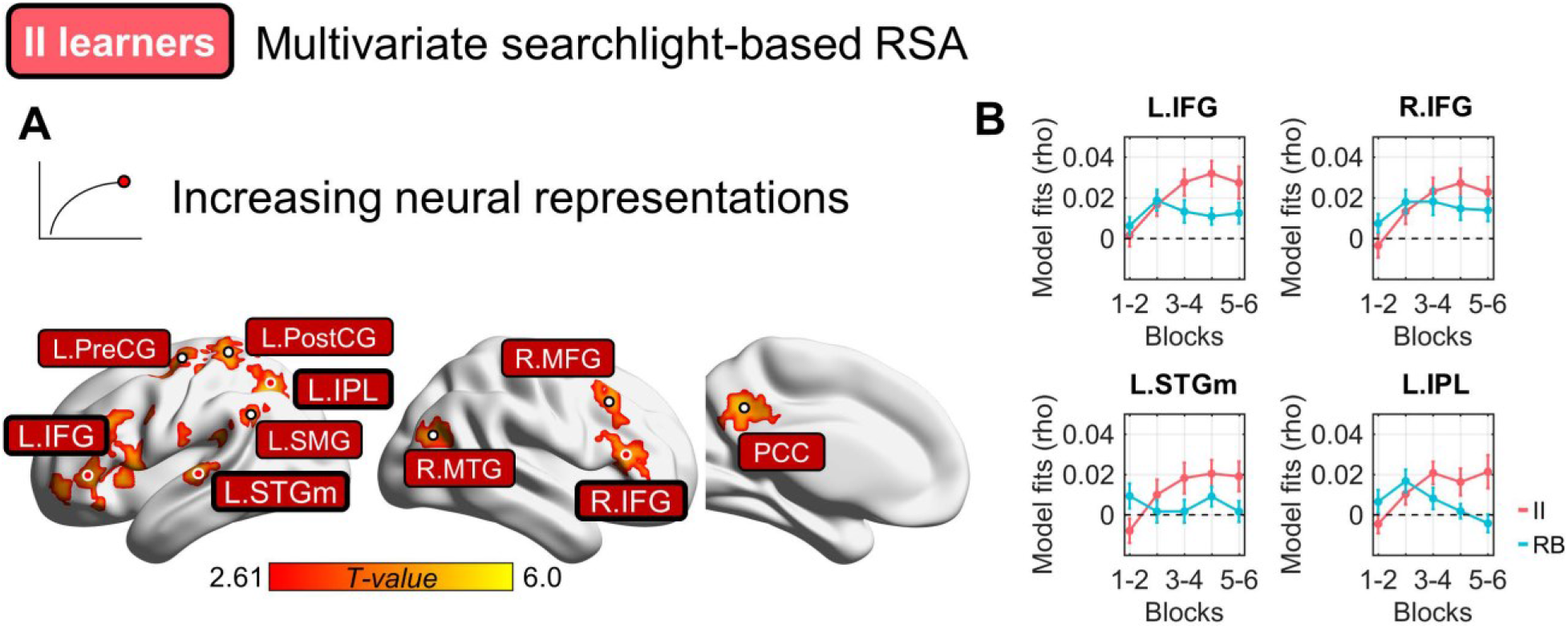
Changes in multivoxel representations of category distance during learning for the II learners. **A**, a fronto-temporoparietal network showed significantly increasing neural representations of category distance for the II group. No significantly decreasing or inverted-U pattern was found. Brain maps were thresholded at the voxel-level *P* = 0.005 and cluster-level FWER = 0.05. **B**, *post hoc* ROI-based RSA analysis for four representative ROIs (highlighted in panel A). The RSA model fits for the RB learners were also displayed for the same regions for visual comparison purposes.

**Table 1:** Brain clusters that showed different emerging patterns of the neural representations of category distance.

| Group \<br>pattern | Regions | Peak MNI<br>coordinates (mm) |  |  | Cluster size<br>(# voxel) | Peak t value |
| --- | --- | --- | --- | --- | --- | --- |
|  |  | x | y | z |  |  |
| **II** Increase | L.IFG | -54 | 35 | 7 | 512 | 4.14 |
|  | L.PreCG | -30 | -1 | 64 | 271 | 4.75 |
|  | L.SMG/IPL | -39 | -40 | 13 | 123 | 4.09 |
|  | L.STGm | -60 | -19 | 7 | 96 | 3.91 |
|  | L.Precuneus<br>/IPL | -12 | -55 | 40 | 512 | 5.04 |
|  | R.IFG/MFG | 39 | 20 | 43 | 313 | 3.75 |
|  | R.MTG | 48 | -76 | 22 | 104 | 4.53 |
| **RB** Increase | SMA | -6 | 32 | 43 | 306 | 4.25 |
|  | R.IPL | 36 | -37 | 43 | 131 | 4.22 |
|  | R.PreCGd | 12 | -10 | 67 | 205 | 4.39 |
| Decrease | L.STGp | -60 | -37 | 13 | 225 | 4.46 |
|  | L.MOG | -30 | -85 | 13 | 140 | 3.99 |
|  | R.STG | 63 | -34 | 19 | 83 | 3.71 |
| Inverted-U | R.PreCGv | 54 | -1 | 49 | 277 | 4.51 |
|  | R.IPL/SPL | 39 | -52 | 64 | 140 | 4.45 |
|  | R.ACC | 6 | 20 | 25 | 100 | 3.94 |
|  | R.Precuneus | 9 | -70 | 61 | 572 | 4.85 |
|  | R.Hipp/<br>Parahipp | 24 | -22 | -17 | 192 | 4.59 |

### RB learners: The emerging neural representations of category-perceptual distance

For the RB learners, univariate activations related to sound categorization were similar to that of II learners (Fig. S2C, SI). The decreasing activations were restricted to the dorsal medial prefrontal cortex, supramarginal gyrus (SMG), and middle temporal regions (Fig. S2D, SI). No region with other changing patterns was found. Importantly, for the multivariate neural representations, we identified regions showing three types of dynamic patterns. For the decreasing pattern, we identified three regions, including the bilateral STG and the left middle occipital cortex (L.MOG) (Fig. 6A). Intriguingly, we also identified extended brain areas showing the inverted-U pattern, including the R.PreCG, R.IPL/SPL, right inferior ACC (iACC), right precuneus, right hippocampus (R.Hipp), and parahippocampus (Fig. 6B). This finding indicates that these regions encode category-perceptual distance information prominently in the middle of the training (i.e., block 3-4) and the representations decreased at the end of the training. For the increasing pattern, we identified three significant regions (Fig. 6C), including the SMA, right dorsal PreCG, and IPL. These dynamic emerging patterns for the RB learners were schematically summarized in Fig. 6D (see Table 1 for detailed regional statistics). In summary, the bilateral STG shows the representation at the early phase of training and decreases afterward while the right frontoparietal regions, ACC, and hippocampus encode the category information dominantly in the middle of training. The right IPL, PreCG, and SMA represent the category information dominantly in the late phase of training. These dynamic patterns indicate that RB learners’ neural representations did not emerge linearly in specific regions, instead, it might be gone through a transformation in representation across the above regions during training.

**Fig. 6.**
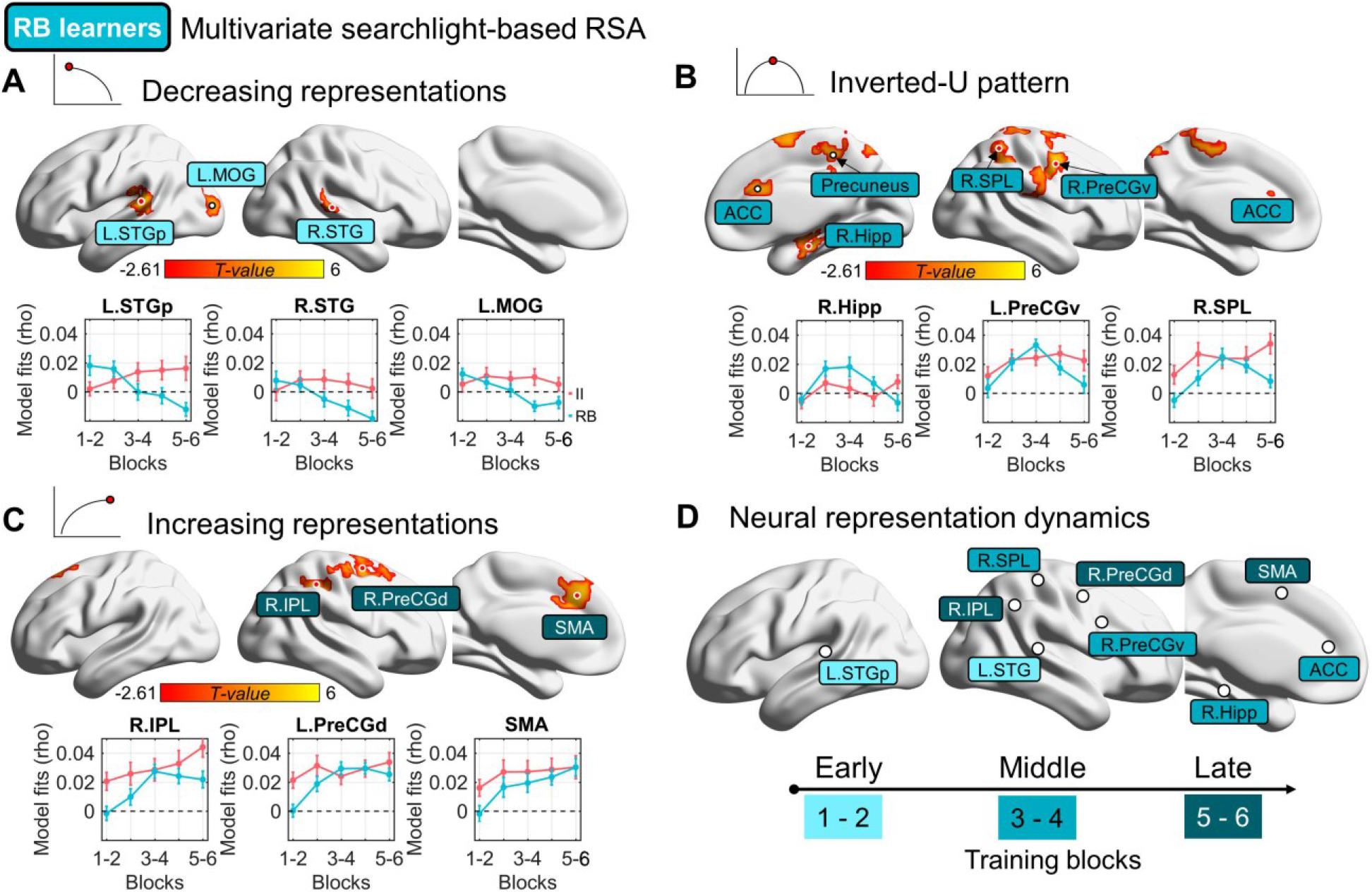
Dynamic changing patterns in multivoxel representations of category distance for RB learners. **A**, decreasing neural representations in the bilateral STG and left middle occipital gyrus. Line graphs on the lower panel show the changing patterns across training blocks. The neural representations for the II learners were also shown for visual comparison in the same region. **B**, the inverted-U pattern in the neural representations. The neural representations in the frontoparietal regions, anterior ACC, and hippocampus emerged prominently in the middle of training. **C**, increasing neural representations were found in the right IPL, dorsal PreCG, and SMA. All the searchlight maps were thresholded with voxel-level *P* = 0.005 and cluster-level FWER = 0.05. **D**, a schematic summary of the spatiotemporal representational dynamics. The region positions are derived proximately from the A-C panels (the peak coordinates). The lightness denotes three pairs of training blocks (i.e., early [block 1-2], middle [block 3-4], and late [block 5-6]). ROI abbreviation: L/R.STG, left/right superior temporal gyrus; L.MOG, left middle occipital gyrus; SMA, supplementary motor area; R.Hipp, right hippocampus; L.STGp, left posterior portion of superior temporal gyrus; R.IPL, right inferior parietal lobule; R.SPL, right superior parietal lobule; ACC, the middle section of the anterior cingulate cortex; R.PreCGd, right dorsal precentral gyrus; R.PreCGv, right ventral precentral gyrus.

## Discussion

We examined the neural dynamics underlying the acquisition of two different types of auditory category structures using a feedback-based sound-to-category training paradigm with fMRI and multivariate pattern analyses. The same underlying dimensions, spectral and temporal modulations, considered the building blocks of speech and music, were common to both category structures. For the RB category structure, optimal performance during learning is obtained by selectively attending to the spectral and temporal modulation dimensions and developing and validating rules that map sounds to categories. In contrast, the II task required pre-decisional integration of the spectral and temporal modulation dimensions by leveraging procedural-based learning of auditory-motor pairings through reinforcement. The feedback-based training procedure was identical for the RB and II tasks. Our design allowed the test of competing models: do representations of the auditory dimensions emerge within a processing hierarchy (the STG-IFG pathway) that are functionally specialized in mapping acoustic signals to behaviorally relevant categories irrelevant of stimulus structure? Or do the emerging representations reflect the task at hand, varying dynamically based on the category structure and underlying learning strategies? Our results point to the latter. Emerging representations do not follow a strict hierarchy and the extent to which representations emerge within the auditory-motor regions versus regions involved in cognitive control are determined by the dynamics of the category learning task (RB vs. II).

Despite large individual differences in learning success, participants learned RB and II tasks to similar extents. Notably, though, they tended to use different decisional strategies, with the II task yielding more procedural-based strategies, and the RB task yielding more rule-based strategies. Importantly, corrective feedback was also differentially processed across tasks as a function of the time course of learning within a corticostriatal network. The putamen and caudate nucleus are increasingly sensitive to feedback processing in the RB task, whereas distributed fronto-temporoparietal regions showed decreased activations in response to feedback for II learners. Representational similarity analysis (RSA) revealed emerging representations of category distance in a bilateral frontoparietal network for both groups. A dorsal sensor-motor pathway, especially the left superior temporal gyrus (STG) and left triangular section of the inferior frontal gyrus (IFGtri) both show increasing representations over training blocks in the II task. These emerging neural representations relate to both training-induced behavioral improvements and individual differences in learning success. In contrast, the category-distance representations in this dorsal pathway *decreased* over training blocks in the RB task. This suggests that representations within these regions are not key to the RB learning task; instead, emerging representations within a large, distributed frontal-parietal-hippocampus network involved in cognitive control are key to learning RB categories. These novel findings are consistent with the dual-learning systems model for auditory category learning and detail commonality and differences in the dynamics of multivoxel neural representations and feedback-related cortico-striatal involvement as a function of the category structures are learned.

For the II structure, pre-decisional integration and procedural-based learning are key for optimal learning. We found a dorsal auditory-motor pathway consisting of the left superior temporal gyrus (STG), supramarginal gyrus (SMG), anterior inferior parietal lobule (IPLa), and precentral gyrus (PreCG) shows increasing representations of perceptual category distance. This auditory-category representational network is consistent with a broad speech processing network, spanning the frontal, temporal, and parietal lobes (Feng et al., 2021; Giraud and Poeppel, 2012; Hickok and Poeppel, 2007; Rauschecker and Scott, 2009). Among these brain regions, the STG has been demonstrated to encode multidimensional acoustic signals (including spectral and temporal modulation) that differentiate native speech categories (Arsenault and Buchsbaum, 2015; Bonte et al., 2014; Feng et al., 2016; Feng et al., 2018; Formisano et al., 2008; Mesgarani et al., 2014), which is presumably acquired slowly with the development of one’s native language. Our findings, together with previous observations that short-term training on non-native speech and non-speech auditory categories are associated with increased neural recruitment of the STG, suggest that the STG plasticity occurs rapidly for adult speech learners (Callan et al., 2003; Desai et al., 2008; Feng et al., 2019; Leech et al., 2009; Wang et al., 2003; Zhang et al., 2009). We posit that multivoxel representations in the STG may increasingly encode perceptual boundaries among the four categories of the II structure, but not RB structure, by increasing the sensitivity of between-category sounds while decreasing the sensitivity of the within-category items. We do not argue that these emergent perceptual-distance neural representations are abstract or categorical, instead, these emergent representations are sensitive to perceptual similarities between sounds. We speculate that the neural representations could transform to abstract or categorical with extensive training (Reetzke et al., 2018). Besides the STG, we identified the triangular section of the left inferior frontal gyrus (L.IFGtri) showing the category-structure-specific emerging representations over training sessions. This L.IFGtri is located within the ventral IFG. Previous studies have found that the IFG is involved in the acquisition of auditory or non-native speech categories (Lee et al., 2012; Myers, 2014; Myers and Swan, 2012) with response patterns in this region showing greater sensitivity to between-category changes and reduced sensitivity to within-category changes after training. To achieve successful learning, the IFG is hypothesized to work dynamically with the temporal cortices (e.g., STG) to form categorical representations and make overt categorization decisions (Feng et al., 2021; Myers, 2014; Myers et al., 2009; Myers and Swan, 2012). Here we demonstrate that different sub-regions of the IFG showed distinct profiles of plasticity. While the L.IFGtri shows increasing representations for the II learners but a plateau-like representation dynamic for the RB learners (see Fig. 4B and Fig. 5B), the lateral IFG yielded similar increasing representations for both groups (Fig. 4A). These findings suggest that the L.IFGtri may be related to the acquisition of task-specific (procedural-based) category structures, while the lateral IFG may be related to *task-general* sound-to-category learning and decisions.

In contrast to the unidirectional increasing representation for the II task, we found a complex dynamic pattern in neural representations for the RB task, including decreasing, inverted-U, and increasing dynamics. First, decreasing representations of perceptual category distance are found in the bilateral STG, which suggests that the neural sensitivity to the perceptual similarity between sounds was reduced as a function of training. This finding suggests that as learners use more rule-based learning strategies, categorization decisions are less driven by bottom-up perceptual information. Instead, the emergence of higher-order abstract rules may guide categorization decisions. Second, we found a distributed frontal-medial temporal network show an inverted-U emerging pattern, in which this network represents the perceptual information highest in the middle of training while decreasing at the late phase. This network consists of frontoparietal regions, hippocampus, and anterior cingulate cortex (ACC) that have been associated with functions of working memory, knowledge-based learning, and memory formation. The frontoparietal regions may support the encoding of stimulus-derived perceptual distance for rule generation, testing, and maintenance that are highly related to working memory and executive controls. This emergent representational network is consistent with predictions from multiple-learning systems models (e.g., Competition between Verbal and Implicit Systems (COVIS) (Ashby et al., 1998) and dual learning system (DLS) model (Chandrasekaran et al., 2014a; Chandrasekaran et al., 2014b). The frontoparietal regions overlap with the executive brain network includes the bilateral prefrontal cortices, IPL, and ACC. These regions have been proposed to be integral to hypothesis-testing (Ashby et al., 1998; Ashby and Maddox, 2005; Ashby and Valentin, 2017; Chandrasekaran et al., 2014b). The frontoparietal representations during learning may be in support of domain-general working memory and executive control processes, which enable the transformation of the perceptual information into testable rules that are stored temporally in the hippocampus (Radulescu et al., 2019). Lastly, the increasing pattern of neural representations of category distance was located in the right dorsal motor system, including the right parietal, precentral, and supplementary motor areas (SMA). The neural representations in these regions are more categorical, which suggests that the regions may support the emerging representation of rules or rule related category decisions. Altogether, we demonstrate that the neural representations of category distance for the RB learners undergo dynamic transformation across training, where decreasing dependence of perceptual information in the STG, temporally representing perceptual information in a frontoparietal-medial temporal network, and increasing representation in the motor network.

## Summary and conclusion

Using decision-bound computational modeling, univariate activation, and multivariate representational similarity analyses, we demonstrate partially dissociated neural dynamic patterns in response to learning information-integration (II) and rule-based (RB) category structures. A sensory-motor pathway, prominently in the superior temporal and inferior frontal regions, and dorsal precentral gyrus showed increasing multivariate representations of category information during the process of learning II categories, while a frontoparietal-hippocampus network showed a complex dynamic representational pattern for learning RB categories. Moreover, univariate analysis showed that feedback-related activations of a frontostriatal network increased for RB learners whereas it decreased for II learners. These findings demonstrate that learners’ corticostriatal systems are highly plastic and sensitive to the types and dynamics of the category learning task. Emerging auditory category-related representations are not strictly restricted to regions (e.g., STG) that are functionally specialized to process key perceptual dimensions. Instead, consistent with recent neural models, learning to categorize is flexibly achieved by strategically learning efficient and task-dependent representations (task-state representations). The composition of category structures, and consequently the dynamics of decisional strategies (sound-to-rule vs. sound-to-reward mapping) during the process of learning are key determinants in the emergent representational dynamics underlying auditory category learning.

## Acknowledgments

This work was supported by grants from the National Institute on Deafness and other Communication Disorders of the National Institutes of Health under Award No. (R01DC013315 834 to B.C.) and the General Research Fund (Ref. No. 14619518 to G.F.) by the Research Grants Council of Hong Kong.

## Supplementary Information

**Fig. S1.**
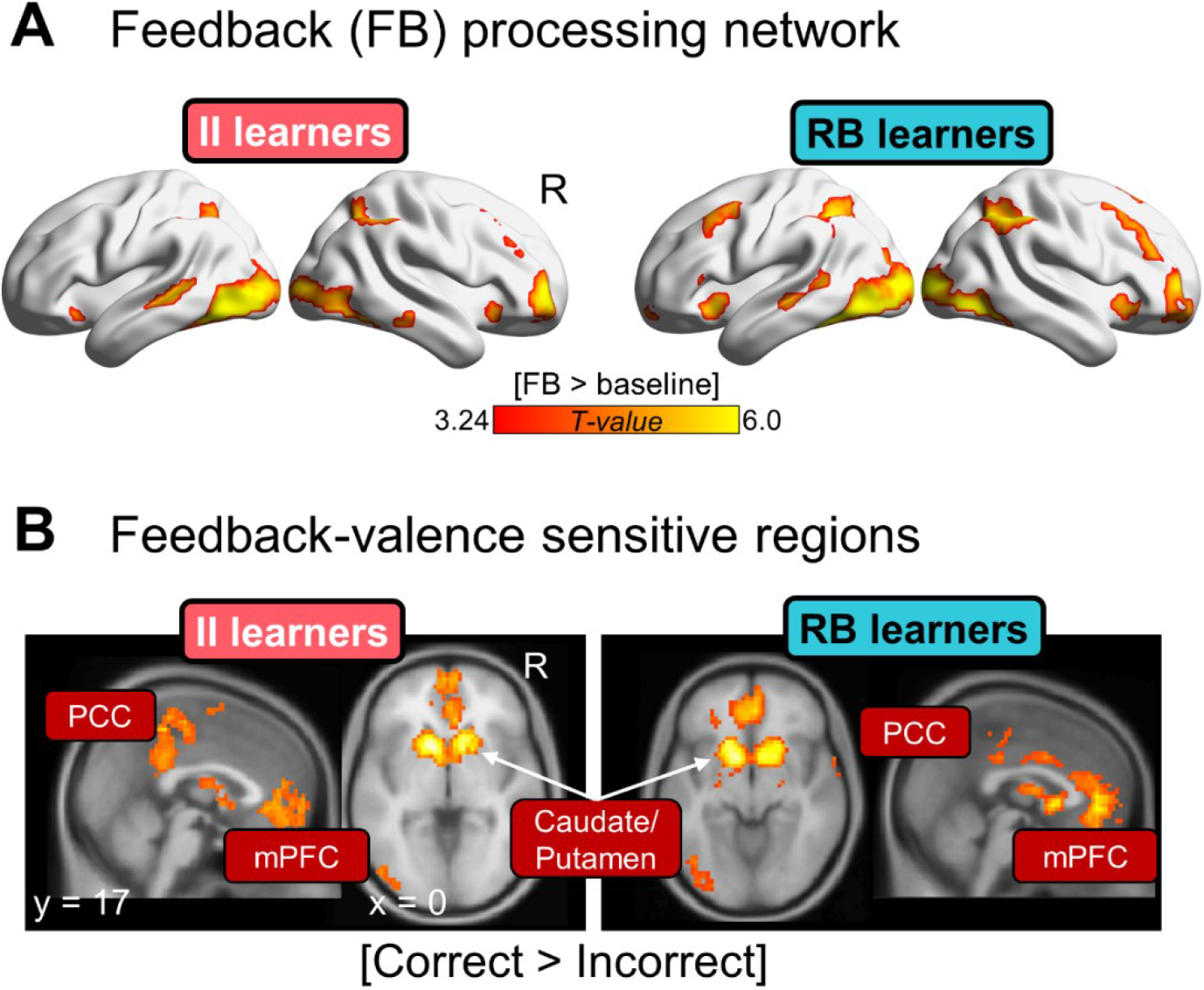
Univariate feedback-processing-related activations for each learner group. **A**, Feedback (FB) processing brain networks (FB vs. baseline) in each group. Activations were comparable between the two groups. No significant group difference was found (collapsed across blocks). Voxel-level *P* = 0.001 and cluster-level FWER = 0.05. **B**, corticostriatal regions that are comparably sensitive to correct feedback (correct vs. incorrect feedback) between the two groups. No significant group difference was found (collapsed across blocks). Voxel-level *P* = 0.001 and cluster-level FWER = 0.05.

**Fig. S2.**
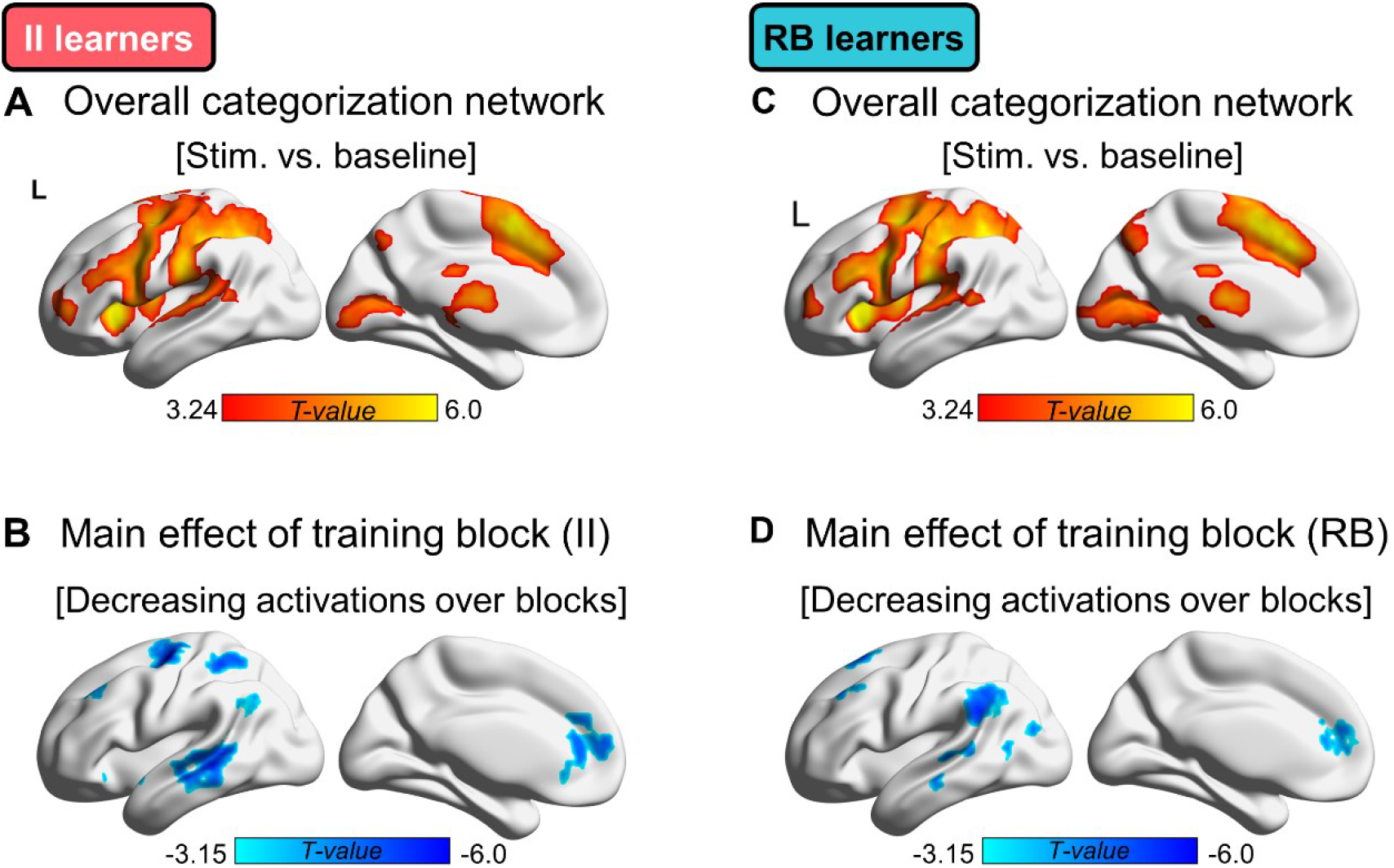
The sound-categorization-related brain activations and the activation changes over training blocks. **A**, a distributed fronto-temporoparietal-subcortical network was activated during sound categorization for the II learners. **B**, decreasing categorization-related activations following training in the dorsal prefrontal, precentral, inferior parietal, angular, anterior cingulate, and middle temporal regions. No region was found showing a significant increasing or inverted-U pattern. **C**, a similar distributed fronto-temporoparietal-subcortical sound-categorization network was found for the RB learners compared to the II learners. **D**, decreasing categorization-related activations were found prominently in the temporoparietal, prefrontal, and anterior cingulate areas. All the above activation maps were thresholded at the voxel-level *P* = 0.001 and cluster-level FWER = 0.05.

